# Advancing FAIR Data Management through AI-Assisted Curation of Morphological Data Matrices

**DOI:** 10.1101/2025.07.08.663621

**Authors:** Shreya Jariwala, Brooke L. Long-Fox, Tanya Z. Berardini

## Abstract

Curation of biological and paleontological datasets is a labor-intensive process that requires standardization and validation to ensure data integrity. In particular, manual curation of datasets is prone to human errors such as typographical errors, inconsistent formatting, and incomplete metadata, which hinder reproducibility and compliance with Findability, Accessibility, Interoperability, and Reusability (FAIR) principles. Artificial Intelligence (AI) offers a transformative solution for enhancing research efficiency by automating data validation, improving accuracy, and streamlining curation workflows. This study presents a proof-of-concept implementation of an AI-assisted curation tool developed for MorphoBank, an open access repository established to enhance standardization and usability of morphological character datasets. Specifically, this work presents an AI tool designed to extract, structure, and standardize morphological character data from published literature into the NEXUS file format, a widely used format for phylogenetic analyses. This tool leverages machine learning techniques, including Large Language Models (LLMs), to automate the extraction of character names and states from text in various formats, reducing manual data entry errors and improving data completeness. The system enables efficient conversion of matrix-only files into complete, machine- and human-readable datasets that include key character metadata. By assisting with these tasks, the tool reduces the manual effort required for curation while improving consistency and standardization. This approach increases the FAIRness of morphological character data and provides a framework for extending AI-assisted curation to other types of biological data. These results illustrate the potential of AI-assisted workflows to support scalable data curation and reuse in paleontology, systematics, and evolutionary biology.

## Introduction

Artificial intelligence (AI) is transforming scientific research across disciplines, including how data are curated, validated, and reused. Large Language Models (LLMs) and other AI-powered tools are already assisting with tasks such as drafting scientific manuscripts, generating code, and annotating data [1–4]. While AI has long been applied in areas such as drug discovery and genomics [5, 6], its application to evolutionary biology and ecology (such as automated detection of morphometric landmarks and the conceptualization of AI-based digital curators for biological collections) is more recent [7–9]. These emerging approaches suggest a critical role for AI in supporting scalable and standardized curation of complex datasets in paleontology, biology, and other scientific fields.

This study builds on these approaches by implementing an AI-assisted pipeline for curating morphological character matrices from published literature, designed to facilitate reuse in open-access repositories, such as MorphoBank (www.morphobank.org). The system leverages Machine Learning (ML) models to extract morphological character names and character states from scientific publications and integrates the character information into a structured NEXUS file format for phylogenetic analyses. From a computational perspective, this task can be framed as structured information extraction from semi-structured scientific documents, a problem that remains challenging due to heterogeneous formatting, complex tables, and domain-specific terminology. By reducing the time and effort required and improving accuracy and standardization, the developed tool offers a powerful solution for increasing the FAIRness of morphological datasets.

Morphological datasets are foundational to biology and paleontology, encoding traits (characters) and their observed variation (character states) across taxa, and are often represented as a matrix with taxa in rows and characters in columns. Each matrix cell stores the coded state for a given taxon–character combination (with conventions for polymorphism, uncertainty, and missing data depending on the dataset). However, morphological character matrix curation is a time-intensive and error-prone process [10–12]. Many morphological character matrices exist only within published literature, often embedded as tables within the paper. Researchers must often manually extract character descriptions from publications, standardize terminology, and reformat data into structures files such as NEXUS [13] or TNT. Inconsistencies in character definitions, formatting, and missing data make this task particularly challenging and susceptible to transcription errors. Scalable, automated curation tools are urgently needed to close these gaps.

The novel curation tool presented here illustrates AI’s potential to close the gap in reliable, robust, automated data curation technologies by supporting large-scale paleontological and biological data curation and offering a framework for expanding AI-assisted curation approaches to other subdisciplines. This AI-assisted curation tool is currently implemented at MorphoBank, a digital repository for morphological data that facilitates real-time collaboration and dataset archiving. In collaboration with the Paleobiology Database (PBDB) [14], 595 NEXUS files were transferred to MorphoBank. However, these files lacked critical metadata (including the vital and curation effort-intensive CHARACTER blocks), making the data difficult to interpret and reuse without referring to the original publications. Initial data transfer efforts relied on manual data curation which was a labor-intensive task due to the highly variable structure, number, and length of character names and states. On average, a 100-character matrix took upwards of two hours to curate manually. The AI-assisted curation tool streamlines the extraction of morphological character name and state data from published articles and merges it with the matrix-only NEXUS file to create a final file with complete character information.

To increase efficiency in adding character names and character states to the NEXUS files, a web-based AI tool was developed to automate the extraction and integration of character information to NEXUS files for deposition as FAIR data in MorphoBank. The AI-assisted curation tool extracts morphological character name and state data from published articles and merges it with the matrix-only NEXUS file to create a final file with complete character information. This implementation demonstrates the utility of AI for large-scale data curation and sets a precedent for extending similar tools to other biological data types and repositories. The goal of this study is not to present a finalized production system but rather a proof-of-concept demonstration that LLMs can assist in extracting morphological character information from published literature and structuring it for data reuse. The implementation described here was developed to support MorphoBank curation workflows and to evaluate whether AI-assisted extraction can substantially reduce manual effort required to reconstruct character metadata from legacy publications.

## Materials and Methods

This study developed a proof-of-concept AI-assisted curation tool, called MatrixCurator, to automate the extraction and structuring of morphological character data from scientific literature into the community-standard NEXUS file format. The tool uses an LLM and a modular pipeline that includes preprocessing, LLM-assisted data retrieval and validation, structured schema conversion, and performance evaluation (Fig. 1). A central methodological feature of the system is a multi-agent architecture in which a retrieval agent generates candidate character-state extractions and an independent evaluation agent verifies them against the source text, enabling iterative correction before final data serialization. The workflow begins with user inputs: a file (e.g., .docx, .pdf) representing part or all of the journal article or supplemental data, a .nex file that contains only the data matrix and taxon information, character count specifications, the relevant page range, and the selection of an appropriate LLM document parser. The relevant pages of the article are then spliced and parsed for data extraction. A Retriever Agent extracts the characters data as a JSON (JavaScript Object Notation) object. The structured data, now a JSON object, undergo two validation steps: 1) a Count Evaluator that checks if the extracted content adheres to character count specifications, and 2) a Data Evaluator that verifies the accuracy of the extracted data. If either validation step fails, the process iterates back to the Retriever Agent for re-extraction and re-structuring. Upon successful validation by both evaluators, the JSON schema is converted into the final, formatted NEXUS (.nex) file. The pipeline is implemented as a web-based app and has been integrated into workflows for morphological character matrix curation in MorphoBank.

**Figure 1:**
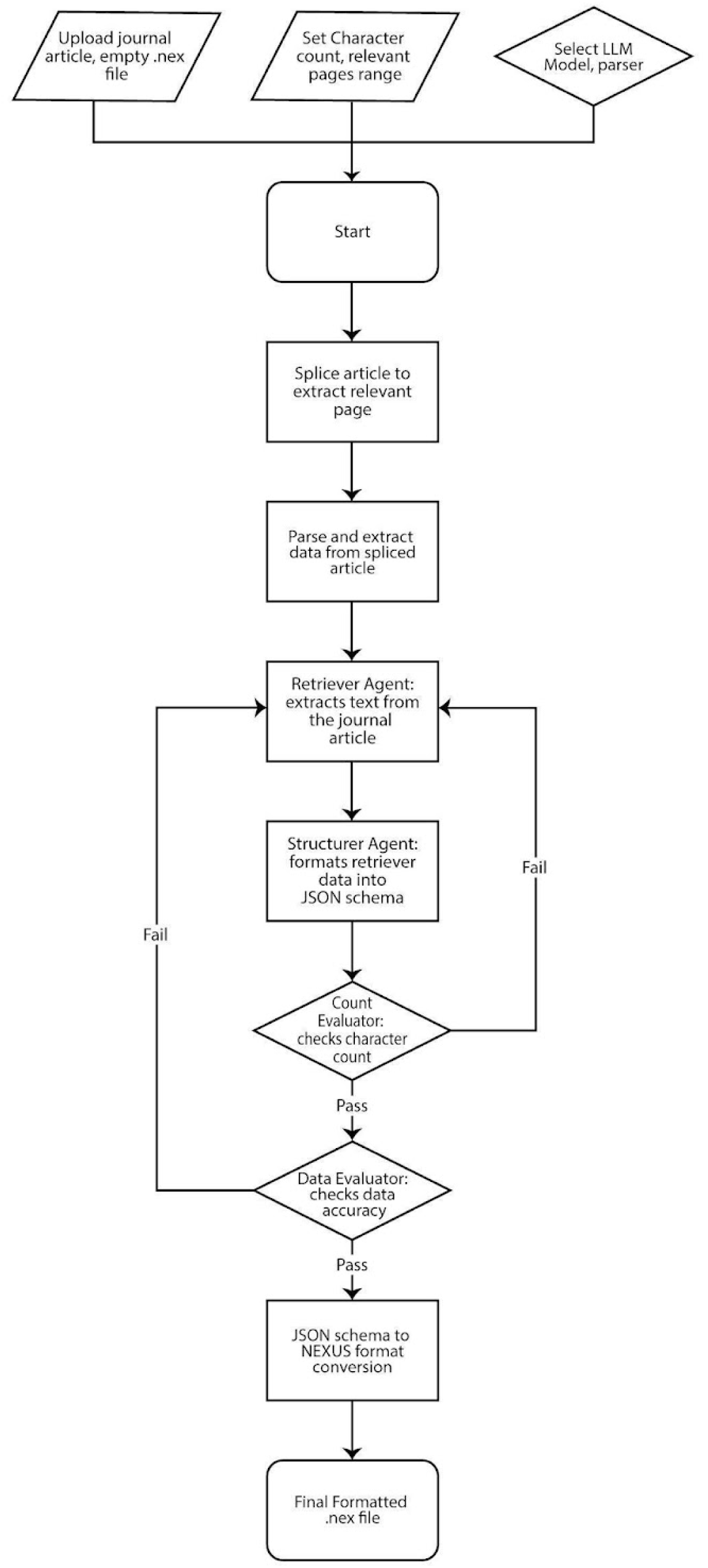
This Large Language Model (LLM)-assisted pipeline automates the conversion of journal articles into the NEXUS (.nex) format through a multi-step process.

### NEXUS File Format

The NEXUS file format is a standard widely used in systematics and phylogenetics to encode different types of data [14]. It consists of structured blocks of data including (but not limited to) TAXA, CHARACTERS, MATRIX, and TREES. The TAXA block lists the organisms used in the research, while the CHARACTERS block defines the morphological traits or features being compared amongst the organisms in the TAXA block. The MATRIX block contains the character state scores for each taxon in the dataset. These values are essential for phylogenetic inference.

While not all blocks are required to run an analysis, the CHARACTERS and MATRIX blocks are essential for morphological character data analyses; they encode both the structure and content of the dataset. An optional but highly useful feature, the STATELABELS command within the CHARACTERS block, allows researchers to define each character and the character states. This feature is critical for making morphological datasets FAIR, as well as providing reusable data to enable replicable, reproducible, and transparent research. The STATELABELS command lists each character by number, followed by a label describing the character, and, if desired, a list of possible states for that character. The AI-assisted pipeline described in this paper was developed to reconstruct these missing blocks by extracting and standardizing the character descriptions from the original publications. In the example from [15] shown in Box 1, STATELABELS defines the meaning of each binary-coded character and the MATRIX block encodes which state (0 or 1) each taxon possesses. This structure of the NEXUS file is both human- and machine-readable, and nearly all phylogenetic analysis software available today are set up to ingest and analyze data in this standard format.

#### Box 1.

##### Encoding in NEXUS Format

This truncated example demonstrates how a morphological character matrix from MorphoBank P4896 (http://dx.doi.org/10.7934/P4896) is encoded in NEXUS format.

#NEXUS BEGIN TAXA;

TITLE Taxa; DIMENSIONS NTAX=3; TAXLABELS

Afrolucina_lens Anodonia_alba Armimiltha_disciformis

;

END;

BEGIN CHARACTERS;

TITLE Character_Matrix; DIMENSIONS NCHAR=2;

FORMAT DATATYPE = STANDARD GAP = SYMBOLS = “0 1 2 3”; CHARLABELS

[1] ‘Overall shell shape’

[2] ‘Maximum adult length’

;

STATELABELS 1

‘subcircular’ ‘ovate’ ‘obliquely ovate’ ‘subtrigonal’

, 2

‘<5 mm (very small)’ ‘5-24 mm (small)’

‘25-49 mm (medium)’

‘50-86 mm (large)’ ‘>86 mm (very large)’

;

MATRIX

Afrolucina_lens 01

Anodonia_alba 03

Armimiltha_disciformis 13

;

END;

### Data Acquisition and Preprocessing

This study uses an AI-assisted curation tool to extract morphological character data from scientific publications and add these data to a NEXUS-compliant file for incorporation in a specialized repository. The source data is typically available as tables within the published manuscript or as supplementary information. The character information is often in PDF or DOCX formats which present significant challenges for automated extraction due to inconsistent formatting, where character descriptions and state might be spread across multiple columns and pages. Older documents may consist of scanned images packaged as a PDF. To address these challenges, a robust approach to automated data acquisition and preprocessing was developed and is described as follows.

The preprocessing pipeline begins with a manual inspection of the source document to identify and isolate the page ranges containing relevant morphological character data within each document. Concurrently, the character count within the paper is also manually determined, often starting with an index of 0. At the time of this study, page ranges and character counts were specified manually during preprocessing. Although these steps could in principle be automated, early testing showed that automated page detection significantly reduced extraction accuracy due to variability in how character descriptions are distributed across figures, tables, and text in many publications. Manual specification therefore served as a reliability control during this proof-of-concept evaluation. Ongoing development within MorphoBank is focused on automating this step using document structure detection and table recognition methods. The initial step in the preprocessing pipeline involves splitting the document into individual pages containing the identified relevant data using code written in Python 3.12 [16]. The page ranges are then extracted and saved as a new PDF or DOCX corresponding to the original input file type, operations which are done using the Python package PyPDF2 [17] and the open-source Libre Office [18] suite, respectively. These extracted pages are then independently processed by dedicated parsers. For PDF files, PyMuPDF [19] is utilized to parse the pages in their natural reading order. This parser incorporates Optical Character Recognition (OCR) as a fallback mechanism to handle image-based data, ensuring that information contained in the scanned documents is also extracted. DOCX files are processed using Libre Office [18] to convert them into Markdown format, facilitating downstream parsing. In addition to these community-standard parsing techniques, a generative AI document parser known as LlamaParse [20] is utilized. This approach leverages an LLM accessible via the LlamaCloud [20] Application Programming Interface (API) to extract information from documents. Finally, we utilize Gemini’s [21] Native Vision features, allowing the LLM to leverage its multimodal capabilities to understand and integrate both text and visual elements to process documents. Gemini’s Native Vision capabilities accurately transcribe tables, interpret complex multi-column layouts, understand charts, sketches, diagrams, and handwritten text within documents, and then use this textual and visual information to carry out end-to-end tasks. DOCX files are first converted to PDF using PyPDF2 [17] before being processed by Gemini’s Native Vision.

### Large Language Models

As previously mentioned, this study leverages the Gemini family of LLMs from Google AI, specifically gemini-2.5-flash-lite, gemini-2.5-flash, gemini-2.5-pro, gemma3:27b and llama4:16×17b. Gemini models were chosen for their availability on a pay-per-usage basis and their ability to process large input prompts, offering context windows of 1-2 million tokens depending on the specific model. Tokens are the units of text that LLMs process that are roughly equivalent to words or word parts. A 1-2 million token window allows the model to handle the full text of even very long documents. According to Google, 1 million tokens is equivalent to 1,500 pages of text or 30,000 lines of code. This capability is essential for efficiently processing the complete text of large research papers, especially those containing hundreds of characters or detailed studies that span numerous pages.

Our methodology employs a multi-agent (i.e., a system involving multiple AI entities working collaboratively to complete complex tasks) approach. The two distinct agents in our system are the Retriever, designed to extract information about each character as a JSON object; and the Evaluator, responsible for assessing the quality and accuracy of the extracted data. Due to the faster inference speeds and lower cost, the “flash” Gemini models were selected for use in the Retriever agent. The high-speed requirements of the agent, which involve numerous inferences for each character within a paper, necessitate the efficiency provided by the flash models.

Conversely, the Evaluator agent, requiring more nuanced understanding and reasoning to accurately assess data quality, utilizes the more powerful “pro” models. The pro model refers to a larger, more robust version of the Gemini LLM, fine-tuned for improved accuracy, reasoning, and handling of longer or more nuanced prompts. Each agent operates based on a combination of a System Prompt, which guides the model’s overall behavior, and an Instruction Prompt, which provides specific task instructions, along with data from the research paper being analyzed. To optimize the performance of these prompts, we utilized Vertex AI Prompt Optimizer to automatically refine the system instructions associated with our instruction prompts. The Retriever agent processes the parsed data from the research paper and extracts relevant information specific to each character of interest. The extracted character data is then passed to the Evaluator agent, which assesses the accuracy of the retrieved data by comparing it to the original paper. This Evaluator is designed to assess how accurately the model’s output aligns with a reference standard. The evaluation is conducted by comparing three components: the original input prompt provided to the model, the model’s generated prediction, and a human-verified reference response. Specifically, the Evaluator receives the original prompt input, the LLM’s initial response prediction, and the parsed paper content reference. The Evaluator’s 1-10 correctness score was normalized to a percentage scale for reporting average extraction accuracy (e.g., a score of 9.1 corresponds to 91% accuracy). Reported accuracy values therefore represent the mean normalized correctness score across characters within each benchmark paper. This score is then extracted from the evaluation results and returned for further analysis. This evaluation is performed using LangChain’s Scoring Evaluator, which is configured with the correctness criteria. The Evaluator uses a correctness-based scoring function using a preconfigured tool called “labeled_criteria”, which is loaded with the specified LLM. The full system prompts and evaluation prompts used in this study are available in the public GitHub repository accompanying this manuscript. To enhance efficiency and reduce costs, we implemented the Gemini API’s context caching feature. This allows the parsed research paper content to be passed to the model only once. It then caches the input tokens, and refers to these cached tokens for all subsequent requests related to that data source paper. By avoiding repeated transmission of the full paper content for each character to be read, the number of tokens per character request is significantly reduced and, consequently, the cost of each morphological character read request is lowered.

### Structured Data Schema

To enable systematic computational analysis, the LLM is instructed to generate the data as a JSON using a predefined JSON schema. JSON is a widely adopted, language-independent data interchange format that provides a user-friendly, human-readable structure. Its ubiquity across programming languages and compatibility with modern AI APIs make it an ideal standard for structuring and processing model outputs [22, 23]. We used JSON to represent the states associated with different characters or features. The primary structure consisted of a dictionary where each key, denoted as “Character 1” or “Character 2”, for example, represented a specific character or feature being analyzed. The value associated with each key was an array of strings, with each string representing a distinct ‘state’ or characteristic exhibited by that character. Each character in the JSON schema can have a different number of possible states. For instance, Character 1 may have three states (e.g., “State 0,” “State 1,” and “State 2”), while Character 2 may have only two (e.g., “State 0” and “State 1”). Subsequently, this JSON structure was cast into the NEXUS format (.nex); this conversion was necessary to enable the use of phylogenetic software packages for further analysis. Each character and its associated states was represented using the ‘Enumeration’ keyword, followed by the character name and a forward slash (/) separating the character from the list of its states. The states were then listed as individual strings, delimited by spaces. This structure provides a clear and standardized way to encode character-state relationships. For example, the JSON data {”Character 1”: [”State 0”, “State 1”, “State 2”]} would be represented in the NEXUS format as ‘Enumeration’ ‘Character 1’ / ‘State 0’ ‘State 1’ ‘State 2’. Similarly, the JSON {” Character 2”: [”State 0”, “State 1”, “State 2”]} would be formatted as ‘Enumeration’ ‘Character 2’ / ‘State 0’ ‘State 1’ ‘State 2’. This process ensured that the information extracted by the LLM was appropriately formatted for subsequent phylogenetic analyses, leveraging the broad compatibility and well-defined structure of the NEXUS format.

### Algorithm Optimization

To enhance the accuracy of the character retrieval system, a two-stage evaluation process was implemented. This process consisted of the Evaluator Agent, designed to assess the correctness of retrieved character data, and a Count Evaluator, responsible for verifying the completeness of the character set. The Evaluator Agent directly compared the LLM-retrieved character data against the original data. When the LLM provided an incorrect response, that response was stored, and the LLM was subsequently re-prompted with a negative prompt designed to prevent future iterations of the same error. These negative prompts included explicit corrections paired with firm instructions (e.g., “Do not include anatomical regions not mentioned in the original paper” or “Avoid summarizing character states - list them exactly as shown”). This process aimed to refine the LLM’s retrieval capabilities through iterative learning. Initially, the “flash” model was employed as the Retriever Agent due to its advantage in inference speed and cost-effectiveness. However, after the initial testing phase, it was recognized that the Retriever would benefit from the “pro” model enhanced capabilities in handling more complex retrieval tasks. Complementing the Evaluator Agent was the Count Evaluator routine which performed a simple but crucial function: it counted the total number of characters retrieved by the LLM and compared this count to the user-specified character count. The decision to utilize “flash” models, where applicable, stemmed from their superior inference speed and lower cost, allowing for more time and resource-efficient evaluation cycles. Conversely, “pro” models were strategically deployed for tasks requiring higher complexity, such as the refined retrieval phase within the Evaluator Agent.

### Community Cloud Deployment

The application was deployed to Streamlit Community Cloud. The process involved navigating to the Streamlit user dashboard and clicking “New app,” then pasting the link to the application codebase (https://github.com/tair/matrixcurator). The correct branch was selected, and streamlit_app.py was designated as the main file. Necessary application information, such as API keys, were input via the “Advanced Section.” Finally, initiating the “Deploy” button triggered the build and launch process on Streamlit Cloud, which generated a unique application URL. At present, the deployment is only accessible to internal users and is not publicly available; however, the following section outlines how others can deploy the tool locally using their own API keys.

### Linux Host Deployment

To run the application locally, the repository was cloned from GitHub, and the required Python dependencies were installed using pip install -r requirements.txt. The necessary API keys were added to the .streamlit/secrets.toml file. The application was then launched using the terminal command streamlit run streamlit_app.py. To enable persistent background operation, the command nohup streamlit run streamlit_app.py & was used, allowing the application to run independently of the terminal session.

### Data Extraction Performance and Efficiency

To evaluate document parsing capabilities, the performance of PyMuPDF, Pandoc, LlamaParse, and Gemini Native Vision was benchmarked. This benchmark utilized a curated dataset of 12 research publications (Supporting Information and https://doi.org/10.5281/zenodo.17291372), specifically selected because they contain morphological matrices and represent a range of formatting complexities and layout styles. The benchmark compares a human-generated base markdown file base_md with the text extracted from a PDF/DOCX file result and calculates a similarity score between the two texts based on inserted, deleted, or replaced to make the two sequences identical. The closer the score is to 1.00, the more similar the text inputs are.

### Agent Performance and Efficiency

To measure the performance and efficiency of our multi-agent system, we established a benchmark using a dataset of 32 papers published between 1991 and 2020 (Supporting Information and https://doi.org/10.5281/zenodo.17291372). This collection was curated to represent a range of publication years (1991-2020), document formats (PDF and DOCX), and morphological matrix sizes (from 20 up to 250 characters) commonly encountered in legacy phylogenetic literature. The model’s multimodal capabilities were leveraged to process the article files in PDF and DOCX formats were parsed into plain text using LibreOffice.

We measure the performance and efficiency of our multi-agent system by measuring inference time and cost per character. Table 2 summarizes the average inference time and cost for each agent using different Gemini models and open source models. The open source models were benchmarked on a single NVIDIA A100 80GB GPU, with costs calculated based on an hourly rental rate of $6.25. Notably, the evaluation step was omitted for the open source models, as they consistently returned a perfect value of 10 during testing. Therefore, the reported cost and inference time for these models do not include data from this step.

## Results

Results of the data extraction performance analysis are given as average similarity ratios for each parsing module in Table 1. Gemini Native Vision achieved the highest average similarity ratio (0.86), followed by PyMuPDF (0.66), Pandoc (0.59), and LlamaParse (0.46). Table 2 contains the results of the agent performance analysis, specifically the average inference time and cost for each agent using different models. Table 3 shows the average accuracy and success rate for each agent using different Gemini models and open source models.

**Table 1:** Average similarity ratio between extracted text and a base markdown file for each parsing module across 12 research publications.

| Parser | Similarity Ratio |
| --- | --- |
| LlamaParse | 0.46 |
| PyMuPDF | 0.66 |
| Gemini Native Vision | 0.86 |
| Pandoc | 0.59 |

**Table 2:**
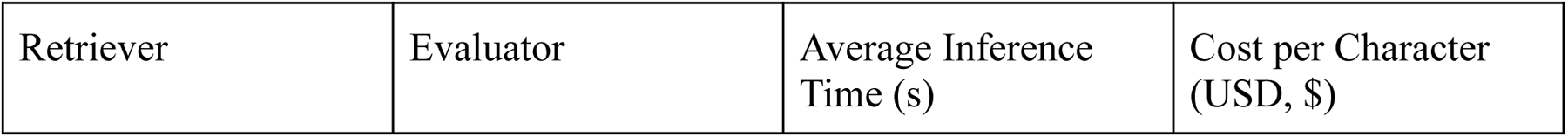
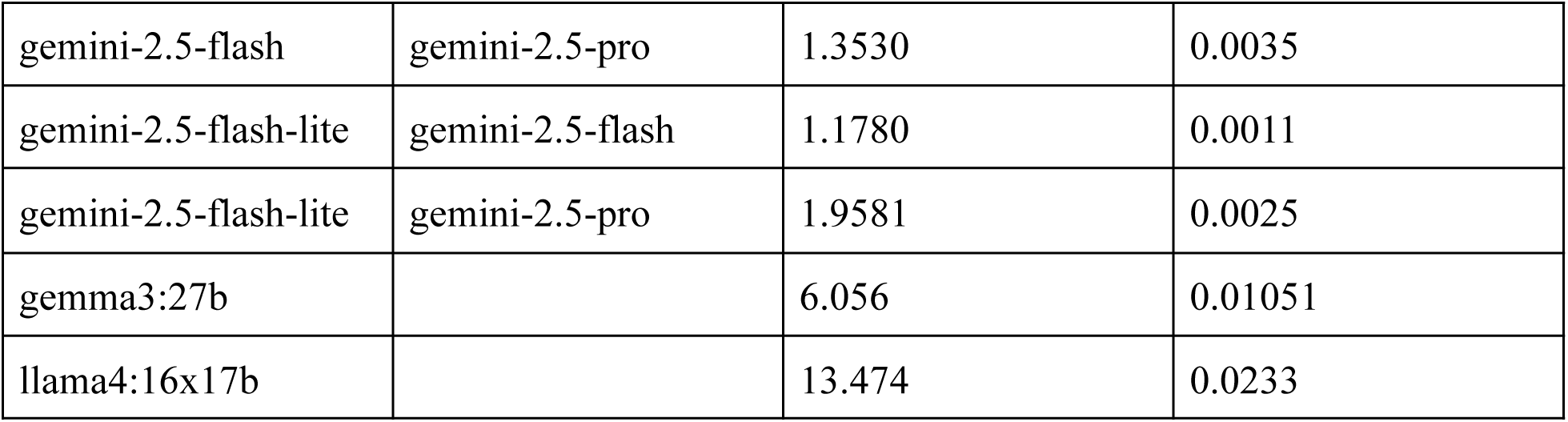
Average inference time and cost for each agent using different Gemini models and open source models.

| Retriever | Evaluator | Average Inference Time (s) | Cost per Character (USD, \$) |
| --- | --- | --- | --- |
| gemini-2.5-flash | gemini-2.5-pro | 1.3530 | 0.0035 |
| gemini-2.5-flash-lite | gemini-2.5-flash | 1.1780 | 0.0011 |
| gemini-2.5-flash-lite | gemini-2.5-pro | 1.9581 | 0.0025 |
| gemma3:27b |  | 6.056 | 0.01051 |
| llama4:16x17b |  | 13.474 | 0.0233 |

**Table 3:** Average accuracy and success rate for each agent using different Gemini models and open source models.

| Extractor | Evaluator | Success Rate (%) | Average Accuracy (%) |
| --- | --- | --- | --- |
| gemini-2.5-flash | gemini-2.5-pro | 99.95 | 90.91 |
| gemini-2.5-flash-lite | gemini-2.5-flash | 96.82 | 85.43 |
| gemini-2.5-flash-lite | gemini-2.5-pro | 95.06 | 85.03 |
| gemma3:27b |  | 99.81 | 30.06 |
| llama4:16x17b |  | 97.54 | 25.09 |

### Computational Efficiency

Leveraging the Gemini API’s context caching mechanism significantly reduced the computational cost of character extraction. To illustrate, consider the analysis of a paper with a matrix containing 164 characters. Without caching, each character request required an average of 1,919 tokens, resulting in a total of 314,716 tokens for all 164 characters. With context caching, an initial 1,820 tokens were used to cache the paper’s content. Subsequent character requests then averaged only 119 tokens each, totaling 19,516 tokens for all characters. This resulted in a combined total of 21,336 tokens with caching, a reduction of approximately 93% compared to the 314,716 tokens required without caching. Assuming a token cost of $0.0000015 per token (based on standard API pricing), the cost without caching would be approximately $0.47, while the cost with caching would be only about $0.03. This dramatic decrease in token usage highlights the practical benefits and cost-effectiveness of employing context caching, especially for large-scale data extraction tasks involving numerous character requests within the same document.

### Curation of Morphological Data

To date, we have applied our AI-assisted curation tool to over 400 published papers containing morphological character matrices (Available at 10.5281/zenodo.17291371). Each publication included a minimum of 20 morphological characters, with some datasets exceeding several hundred characters. In total, the tool has processed over 35,000 character-state entries across a wide taxonomic range. The system was able to extract and structure morphological character data from a range of formatting styles, table structures, and text descriptions encountered in the bench corpus. Despite substantial variation in how characters and states were presented, ranging from fully labeled matrices to minimally described tables, the tool demonstrated consistent performance across the evaluated benchmark dataset in identifying character names, corresponding states, and associated taxa. In curator review, outputs typically required minor edits such as punctuation corrections, normalization of formatting, or alignment of character numbering with the original matrix. More substantial intervention was occasionally needed when source materials were unusually complex or inconsistent. Even in these cases, starting from AI-generated output was considerably faster than beginning transcription from scratch, enabling a substantial reduction in the manual effort needed for dataset preparation. These experiences suggest that the tool can play a valuable role in accelerating morphological data reuse, while still relying on human expertise to ensure accuracy and resolve more challenging cases. By combining automation with curator oversight, the system helps fill a critical gap in supporting large-scale, reproducible phylogenetic research.

## Discussion

This study presents MatrixCurator as a proof-of-concept AI-assisted curation pipeline developed to reduce the manual effort required to recover morphological character metadata from published literature and integrate it into NEXUS files suitable for MorphoBank. Rather than presenting a finished production system, we report a feasibility demonstration: evidence that LLM-based extraction, document parsing, and multi-agent validation can together substantially reduce the manual effort required for a specific, well-defined curation task that has historically bottlenecked progress in making legacy morphological data FAIR-compliant [24,25–27]. The limitations described below are explicitly those of a first-generation proof-of-concept, and several are already being addressed in the next generation of the tool now under active development for direct integration into MorphoBank. The evaluation presented here focuses on demonstrating feasibility within a real-world repository curation workflow rather than establishing a comprehensive natural language processing benchmark. The 32-paper benchmark corpus was curated to represent a range of formatting styles and publication eras commonly encountered in morphological phylogenetic literature. Future work will expand benchmarking to larger and more systematically sampled corpora, incorporate structured error-type logging, and explore automated detection of relevant page ranges and character counts, inputs that are currently provided manually.

### Model and Parser Performance

Model and parser selection proved pivotal to extraction quality.. Among parsers, Gemini Native Vision achieved the highest similarity ratio (0.86) across the 12 document parser benchmark, outperforming PyMuPDF (0.66) and LlamaParse (0.46). This advantage was particularly pronounced for documents containing embedded tables, multi-column layouts, and figures with labeled character states, which were the structures most critical for accurate morphological extraction. Combining high-fidelity document parsing with a multi-agent architecture provided an additional safeguard: the Evaluator agent could identify cases where retrieved content diverged from the parsed source and trigger re-extraction prior to NEXUS conversion. Similar approaches combining LLM-based extraction with validation stages have been effective in other scientific information extraction contexts [28,29]. Among agent configurations evaluated across 32 publications, Gemini 2.5 Pro as Evaluator achieved the highest average accuracy (90.91%) with a success rate of 99.95%. Flash-only configurations produced accuracies in the mid-80% range (85.03-85.43%), reflecting a speed-cost-accuracy trade-off that practitioners can weigh against their curation priorities. These figures represent point estimates from a single benchmark corpus evaluated without confidence intervals or inter-rater reliability assessment, which are recognized limitations for a proof-of-concept study. Future evaluations will incorporate distributional summaries and formal inter-rater analyses. Context caching reduced token use by approximately 93% for a representative 164-character matrix, lowering per-matrix cost from $0.47 to $0.03 and enabling repository-scale processing at feasible cost.

### Open Source Models and Overconfident Outputs

Benchmark analyses also revealed a marked disparity for open-source models. Gemma and Llama reported very high “success rates” (99.81% and 97.54%) yet exhibited low factual accuracy (30.06% and 25.09%), indicating a tendency toward confident but incorrect outputs. They were also slower and more expensive per character than the top-performing Gemini configuration, making them impractical for reliable, scalable deployment in this workflow at present. Limitations remain and motivate curator-led verification. We observed omissions in dense or irregular tables, misalignments when headings repeat, occasional fabricated states, ambiguous anatomical terms resolved incorrectly, and routine normalization issues. Similar challenges have been reported in studies evaluating LLMs for structured information extraction from scientific literature, particularly when documents contain complex layouts, tables, or ambiguous terminology [30,31]. Bias in training data can introduce systematic errors; clear reporting of constraints and ongoing evaluation are essential for trust.

### Limitations and Error Characterization

The system requires human verification at each stage and should not be understood as a fully autonomous curation solution. Automation redistributes curatorial effort from transcription toward verification and quality control rather than eliminating it [9]. This framing is consistent with emerging models of human-AI collaboration in biological collections, in which automated extraction tools support but do not replace expert oversight [9,32–35]. During testing and production use, we identified several recurring error categories that practitioners should anticipate. Table omissions occurred when character rows were missed in dense or irregular tables, particularly where merged cells or complex spanning structures were present. Heading misalignments arose when section headers repeated across pages, causing the Retriever to misattribute character definitions to incorrect sections. State fabrication occurred when source documents described characters qualitatively without enumerating discrete states, prompting the model to generate plausible-sounding but unsupported state labels - a form of hallucination consequential for downstream phylogenetic analyses. Ambiguous anatomical terminology presented additional challenges when the same structure was described inconsistently across publications. Finally, normalization issues such as punctuation and capitalization inconsistencies were common during early testing appeared frequently but were typically resolved with minimal curator effort [29,30]. Because structured edit logging was not implemented during the initial proof-of-concept deployment, frequency estimates for individual error types are not available.

This limitation reflects the exploratory nature of the initial system deployment rather than the absence of quality control during curation.The accuracy figures from the 32-paper benchmark provide the most rigorous quantitative characterization currently available. Future deployments will incorporate structured logging of curator edits by type to support prospective error-rate tracking and targeted improvements to prompts and validation logic. The current workflow requires users to manually specify the relevant page range and character count for each document. Automated scope detection was evaluated in an earlier version of the tool and found to reduce accuracy, particularly for older publications with less structured layouts. Manual input was therefore retained as a deliberate design choice that improves extraction precision and gives curators explicit control over scope. This represents a scaling constraint for high-volume processing, and reliable automation of this step is a primary development goal for the next-generation system currently being integrated directly into MorphoBank – one that will not require manual page range or character count specification.

### User Interface and Workflow Integration

The current prototype is implemented as a Streamlit-based web interface that allows curators to upload documents, specify extraction scope, and review outputs interactively before NEXUS generation. Because the tool is currently deployed internally within MorphoBank workflows, formal usability testing has not yet been conducted. Interface design has focused on enabling curator control over extraction parameters and facilitating iterative review during early-stage development. Within MorphoBank workflows, the interface allows curators to upload source publications, define extraction scope, inspect generated character definitions, and export validated NEXUS files prior to repository ingestion. Future development will include formal usability evaluation, expanded documentation, and deeper integration with MorphoBank’s broader data curation workflows to support wider adoption by researchers and repository curators.

### FAIR Data Compliance

These contributions support FAIR data principles [19]. Findability is improved by converting matrix-only files (which require access to the original publication for interpretation) into NEXUS files with fully labeled CHARSTATELABELS blocks, enabling indexing by trait, taxon, and project within MorphoBank. Accessibility is enhanced because enriched NEXUS files are self-contained and human-readable without reference to external documents. Interoperability is advanced through standardization to NEXUS, which is compatible with widely used phylogenetic analysis tools including TNT, PAUP*, and MrBayes, and through the use of JSON as an intermediate representation compatible with modern AI APIs and downstream data pipelines. Reusability is strengthened by ensuring that character definitions and state labels are explicitly encoded alongside matrix data, providing the context necessary for transparent replication and extension by other researchers [24,25–27].

### Redistribution of Curatorial Efforts

Rather than eliminating human labor, AI-assisted curation redistributes it. Tasks that were historically performed together within individual expert workflows by locating, transcribing or direct copy/paste, and verifying character descriptions, thus become structurally differentiated. Machines draft candidate representations, while human curators retain interpretive authority and responsibility for verification. This redistribution has implications for how biological data infrastructures organize curation workflows and allocate expert effort [9,32–35]. The governance implications of AI-assisted curation workflows are explored in related work currently in preparation. That work examines how structured human quality assurance, including checklist-driven verification and provenance tracking, can support reliable integration of AI-generated outputs into biodiversity repositories. In the proof-of-concept system described here, AI output was explicitly treated as a draft representation, and all generated matrices underwent structured human review prior to repository ingestion.

### Scope, Performance Conditions, and Future Directions

The conditions under which MatrixCurator performs reliably are important for practitioners considering adoption. Performance is strongest for digitally typeset English-language publications in which morphological characters are described in clearly delimited tables or appendices with consistent heading structures and where the document is available in a machine-readable PDF or DOCX format. Performance is expected to degrade when characters are described primarily in narrative prose, when publications are written in languages other than English (which were not evaluated in this study), when documents consist of heavily scanned image-based PDFs where OCR errors compound extraction errors, or when datasets employ highly non-standard or idiosyncratic character description conventions. The current system also does not perform inter-rater reliability assessment or provide performance breakdowns by taxonomic group or publication era; both represent important directions for future evaluation.

More broadly, this work illustrates how artificial intelligence can function within scientific infrastructure as an assistive technology rather than a replacement for expert curation. Automated extraction can accelerate the recovery of structured data from legacy literature, but interpretation, validation, and integration remain human responsibilities [9,33,34]. Effective AI-assisted curation systems therefore depend on hybrid workflows in which automated systems generate draft representations while expert curators retain interpretive authority. Such collaborative approaches may help repositories scale the recovery of legacy biodiversity data while maintaining scientific reliability [25,27,36].

Ongoing development of MatrixCurator focuses on improving extraction accuracy, addressing context-window constraints in large language models, and expanding support for additional dataset formats. Future versions will also integrate the system more directly into MorphoBank workflows and make the tool publicly available to researchers in paleontology, systematics, and evolutionary biology. Although developed for morphological character matrices, the underlying workflow represents a generalizable pattern for AI-assisted scientific data recovery from legacy literature. The combination of document parsing, LLM-based information extraction, and independent validation agents could be adapted to other data-intensive domains in which structured datasets must be reconstructed from heterogeneous publications, including trait databases, ecological observations, and genomic metadata. These developments align with broader efforts to build AI-assisted research infrastructure for biodiversity data. Emerging concepts such as “digital curators” envision AI agents that augment expert practice by supporting tasks such as information extraction, metadata enrichment, anomaly detection, and integration of heterogeneous datasets [37–39]. Within extended digital specimen networks and related biodiversity data infrastructures, such tools may help connect disparate data sources and facilitate large-scale synthesis of biological information [40]. In this context, MatrixCurator represents an early step toward integrating AI-assisted workflows into biodiversity repositories while preserving the central role of expert oversight in ensuring data quality and interpretive accuracy.

## Conclusions

This study demonstrates that AI-assisted workflows can substantially reduce the manual effort for morphological data curation by automating the extraction and structuring of character data from scientific literature and integrating standardized NEXUS files suitable for repository deposition. The MatrixCurator combines document parsing, LLM-assisted extraction, and validation to generate draft character definitions that can be incorporated into MorphoBank.

Importantly, this work represents a proof-of-concept implementation rather than a fully automated curation system. Human expertise remains essential for verifying extracted character definitions, resolving ambiguous terminology, and ensuring consistency with domain-specific standards. By enabling more efficient reconstruction of missing character metadata, this approach supports FAIR principles of findability, accessibility, interoperability, and reusability for morphological datasets. Ongoing development focuses on improving automated scope detection and integrating the system directly into MorphoBank’s curation infrastructure. As repositories increasingly seek to mobilize legacy datasets for reuse, AI-assisted curation workflows may provide an important mechanism for scaling data recovery while preserving the interpretive role of expert curators.

## List of Abbreviations

AI: Artificial Intelligence
FAIR: Findable, Accessible, Interoperable, and Reusable
LLM: Large Language Model
ML: Machine Learning
PBDB: PaleoBiology Database
PDF: Portable Document Format
API: Application Programming Interface
OCR: Optical Character Recognition
JSON: JavaScript Object Notation

## Data Availability

All relevant data are within the manuscript and its Supporting Information files. The MatrixCurator source code and associated prompts are available at GitHub (https://github.com/tair/matrixcurator). Benchmark datasets and project lists used during the evaluation of they system are archived at Zenodo(https://doi.org/10.5281/zenodo.17291372).

## Acknowledgements

This project would not have been possible without the support of the Phoenix Bioinformatics team (in alphabetical by first name order): Alyssa Proia, Amina Khababa, Connie Ng, Eileen Conroy, Erika Bakker, Josh Young, Kartik Khosa, Leonore Reiser, Swapnil Sawant, Trilok Prithvi, and Xingguo Chen. We are grateful for Dr. Maureen A. O’Leary’s continued guidance on MorphoBank. The development and evaluation of the AI-assisted curation tool described here involved the use of multiple LLMs, including Gemini (Google DeepMind), LLaMA the Gemini API and LlamaCloud platforms to assist with document parsing, character-state extraction, and structured output generation. The authors also acknowledge the use of ChatGPT (OpenAI) for language editing, including improving clarity, sentence structure, grammar, and formatting. All intellectual content, research design, and conclusions are the work of the authors. We thank Editor Alexander Dececchi and the four reviewers whose detailed and constructive feedback substantially improved this manuscript by challenging us to think more carefully about the scope and framing of AI-assisted curation workflow, the importance of transparent error characterization, and the governance structures needed to sustain reliable human oversight alongside automated extraction.

